# The Mitotic Chromosome Periphery: A Fluid Coat That Mediates Chromosome Mechanics

**DOI:** 10.1101/2024.12.21.628209

**Authors:** Tania Mendonca, Roman Urban, Kellie Lucken, George Coney, Manlio Tassieri, Neil M. Kad, Amanda J. Wright, Daniel G. Booth

## Abstract

Mitotic chromosomes are specialised packets of condensed genetic material with dynamic mechanical properties. Each chromosome is coated by a sheath of proteins and RNA, called the mitotic chromosome periphery (MCP). The MCP is widely considered as an essential chromosome compartment where its multiple functions bestow material properties important for successful cell division. However, the details of the micromechanical properties of mitotic chromosomes, and specifically if and how the MCP contributes to these features, remain poorly understood. In this study, we present the most comprehensive characterisation of single-chromosome mechanics to date spanning a broadband frequency range, using optical tweezers and a novel microrheology technique. We extend this analysis to the first direct measurements of MCP micromechanics by manipulating levels of Ki-67, the chief organiser of this compartment, and apply a rheological model to isolate its contribution to chromosome dynamics. We report that the MCP governs high-frequency self-reorganisation dynamics and acts as a structural constraint, providing force-damping properties that mitigate mitotic stress. This work significantly advances our understanding of chromosome micromechanics and how the MCP contributes to the fundamental properties of chromosomes.

## Main Text

Chromosomes have intriguing biophysical properties that have been difficult to define despite decades of intense research. Reports on chromosome mechanics vary widely depending on the experimental technique used (1, 2). This variability partly arises from the temporally complex and intrinsically dynamic properties of chromosomes. Depending on the timeframe of the observation window, chromosomes can be described as free polymers diffusing in a viscous nucleoplasm during interphase (3, 4) that then transition to a gel-like (5, 6) state during mitosis. The complex biophysical properties of chromosomes (7), including their emergent behaviours such as condensation, congression and segregation, are influenced by their heterogeneous composition and interactions between the chromatin fibre, proteins, and RNA. While the contribution of some classes of proteins like the structural maintenance of chromosomes (SMC) protein complexes to chromosome mechanics have been well studied (8, 9), the involvement of other chromatin interacting biomolecules remain unclear, or in the case of the mitotic chromosome periphery (MCP), are completely unexplored.

The MCP is a collection of proteins and RNA that redistributes from the disassembled nucleolus to the surface of condensed chromosomes at the onset of mitosis (10–12) and appears to confer biophysical properties to chromosomes, crucial for successful cell division. Chromosomes depleted of the protein Ki-67, the chief organiser of the MCP (13), lack the periphery compartment of over 65 proteins and RNAs (14, 15) and become ‘sticky’ and aggregated (13, 16, 17). Additionally, the MCP appears to be multifunctional, with further roles reported in; promoting chromosome clustering during late mitosis (18, 19), the symmetrical distribution of nucleolar material between daughter cells (13), the maintenance of chromosome architecture, either directly (20) or via organisation of its epigenetic landscape (21), and finally as a protector against DNA damage (22). It has been suggested that some of these functions may be driven by Ki-67 modulating phase separation in chromosomes to keep them “individualised”, ready for segregation (16, 23). Despite the broad range of critical functions attributed to this chromosome compartment, its biophysical properties have yet to be directly tested. To address this gap, we used optical tweezers to perform the first direct measurements of the molecular biophysics of the MCP at the single-chromosome level.

Our **single-chromosome analysis toolkit** includes a stable custom CRISPR-generated Ki-67-mEmerald cell line, enabling direct observation and quantification of Ki-67 on individual human chromosomes. We exploited Ki-67’s role as the chief organiser, to modulate MCP chromosome enrichment, by: 1) using Ki-67 specific siRNA to deplete the MCP and 2) transiently transfecting cells with a custom-designed plasmid gRNA vector as part of a CRISPR Activation system, to increase Ki-67 expression, and therefore enhance MCP recruitment (Figure 1ai and aii, Extended Figure S1 and see methods for details). Chromosomes with wild-type (WT) or altered MCP load (knockdown ‘KD’ or overexpressed ‘OE’) were then isolated from Ki-67-mEmerald cells for single-chromosome analysis using an optical tweezers instrument (C-Trap, LUMICKS) with two independently controlled optical traps, integrated with microfluidics and fluorescence microscopy.

**Fig. 1.**
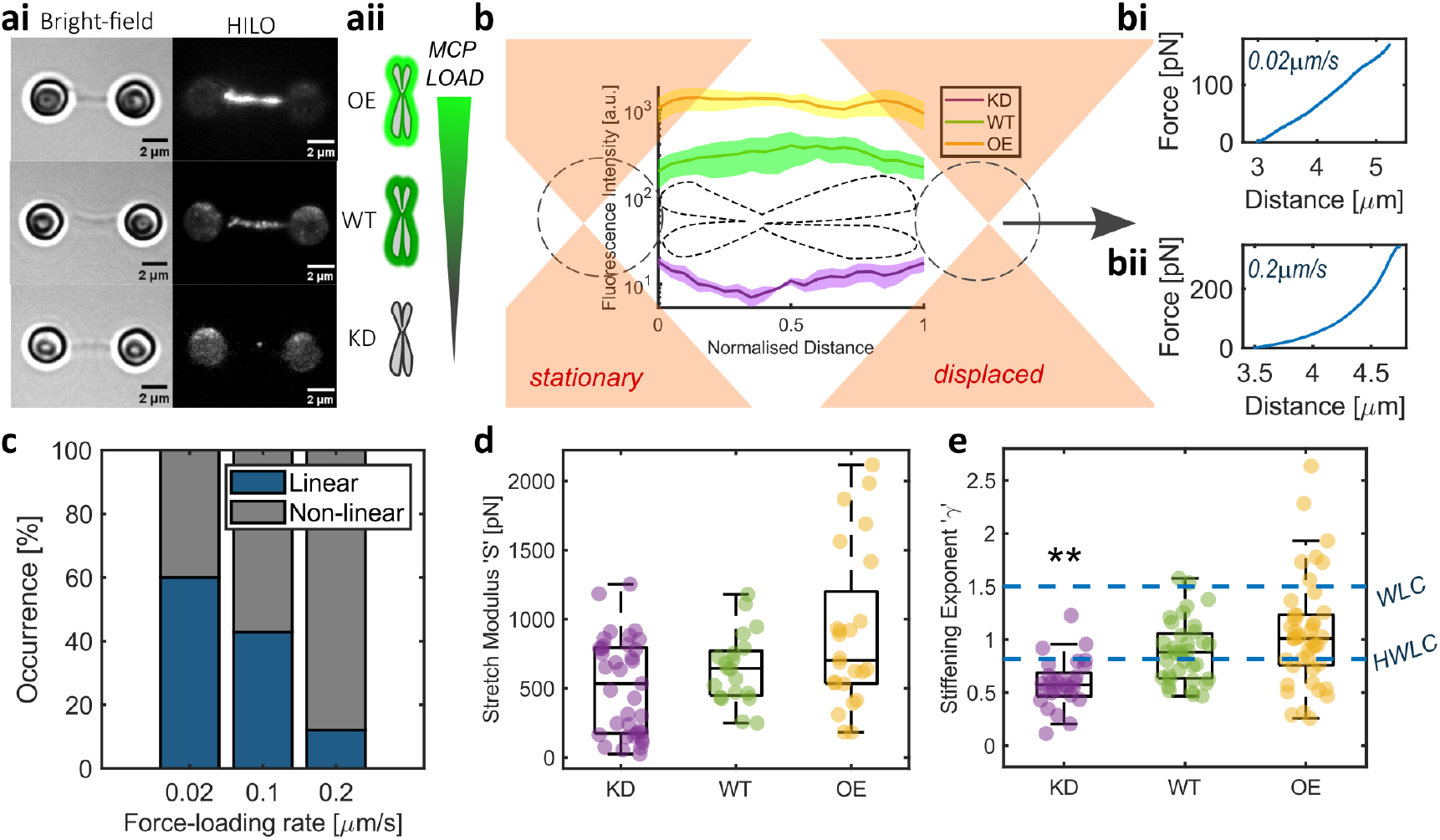
Chromosome mechanics are rate dependent. ai. Examples of Bright-field and Fluorescence HILO images of OE, WT and KD chromosomes in dumbbell configuration. Scale bar = 2 *µ*m. aii. Schematic of MCP load with Ki-67 expression. b. Schematic of the force-extension experiments where one optical trap is displaced while the other is kept stationary, to apply stretching forces at a known speed. Fluorescence intensities of chromosomes were analysed to quantify MCP load. Individual examples of force-extension experiments at either 0.02 *µ*m/s (bi) or 0.2 *µ*m/s rate (bii). showing linear and non-linear behaviour respectively. c. Occurrence of linear and non-linear mechanical response with different force-loading rates in WT chromosomes. d. Stretch modulus was acquired from chromosomes showing linear behaviour at force-extension of 0.02 *µ*m/s. e. Stiffening exponent *γ* from chromosomes showing non-linear behaviour at force-extension of 0.2 *µ*m/s compared to *γ* values for the Worm-like Chain (WLC) and Hierarchical Worm-Like Chain (HWLC) models. Comparisons to WT (Kruskal-Wallis test), significance values: \*\**p* = 0.001

Individual chromosomes were captured between optically trapped pairs of polystyrene bead handles using biotinstreptavidin interactions in a dumbbell configuration (Figure 1ai and b, Methods). A stretching force was applied to each chromosome, by moving the position of one of the laser beams generating the traps which linearly displaced one of the bead handles. Each optical trap functions as a highly sensitive force transducer for small displacements (24), enabling the measurement of picoNewton forces by tracking the displacement of the bead handles relative to the respective optical trap centre, i.e., laser beam focus.

**Chromosomes respond differently to different rates and levels of deformation** (1, 5, 25) with predominantly linear spring-like behaviour when stretched slowly (<0.1 *µ*m/s) or non-linear behaviour in most cases when probed at faster rates (Figure 1bi, bii and c). At a force loading-rate of 0.02 *µ*m/s, mechanical equilibrium is seen in most chromosomes, resulting in a linear increase in force required to stretch the chromosome greater distances (2). For a linear relationship between force and the relative extension of the chromosome, the slope of the graph (e.g. Figure 1bi) defines the stretch modulus ‘*S*’, also referred to as the doubling force (8, 26). This parameter is analogous to Young’s modulus but does not assume material homogeneity. *S* remains unchanged with varying MCP levels (Figure 1d), suggesting the MCP does not influence the elasticity of the mechanically equilibrated mitotic chromosome over long time scales.

At a faster stretching rate of 0.2 *µ*m/s, the force-extension relationship bears a non-linear form in a majority of tested chromosomes (Figure 1bii and c). Such non-linear stiffening can be attributed to the chromosome network microstructure and force transmission through cross-links (28). This stiffening ‘*κ*’ bears a power-law relationship with force ‘*F*’, where *κ* = *aF* ^*γ*^ (27). In WT chromosomes the exponent ‘*γ*’ was 0.88 [0.78 0.98] (mean ± 95% confidence intervals) which is in agreement with literature (27) and lower than the *γ* value of 3/2 attributed to the worm-like chain model used to describe the mechanics of double stranded DNA (27, 29, 30). This decreased stiffening rate has been explained using a Hierarchical Worm-Like Chain (HWLC) model (27) which postulates chromosomes to be hierarchical assemblies of elements with distinct mechanical properties, each acting as a flexible worm-like chain. The *γ* value drops to 0.56 [0.46 0.66] with the loss of the MCP (Figure 1e; Kruskal-Wallis test with multiple comparisons, WT vs KD *p* = 0.001). This lower *γ* value suggests a disruption or loss of structural elements that contribute to force transmission and the sequential stiffening response with increasing force. However, *γ* did not significantly change for OE chromosomes (1 [0.9 1.2]; Kruskal-Wallis test with multiple comparisons, WT vs OE *p* = 0.43) indicating no discernible change to the hierarchical organisation of the chromosome in this case. Individual chromosomes from all treatments exhibited consistent mechanical behaviour with repeated extensions for forces of up to 150 pN, as with previously reported findings (27, 31).

The HWLC model assumes that chromosomes exhibit a predominantly elastic response to stress (27), which contrasts with prior reports of viscous relaxation in chromosomes at force-extension rates of 100 *µ*m/s (25). These results, when considered in isolation, highlight the limitations of discrete single-frequency measurements in accurately capturing the dynamic, time-dependent mechanical properties of chromosomes and the complex contributions of their associated structures such as the MCP.

This prompted us to develop **a novel broadband microrheology approach for single chromosomes**. While micromanipulation tools such as optical tweezers and magnetic tweezers have been used to study interphase chromosomes (3, 32, 33), their potential for broadband characterization of mitotic chromosome dynamics remains underexplored. We have developed a bespoke optical trapping-based stretch-strain method, inspired by a similar approach used for microrheology of cells (34), to extract the mechanics of individual mitotic chromosomes across a broad frequency range. The complex stiffness *κ*^*^(*ω*) derived from this method provides information about the chromosome’s viscoelastic properties. It is defined as the ratio of the Fourier transforms of force *F*(*t*), measured as the picoNewton force exerted by both optical traps on the chromosome with time, and strain *ϵ*(*t*), the relative extension of the chromosome in nanometres over time. *κ*^*^(*ω*) is a complex number with real and imaginary parts that describe the elastic *κ*^*′*^(*ω*) and viscous *κ*^*′′*^(*ω*) components of the chromosome mechanical response.

Using the chromosome dumbbell configuration, chromosomes were stretched by displacing the position of one of the two optical traps a fixed distance of 1 *µ*m, at a rate of 100 *µ*m/s (Figure 2a-b). This force-loading phase was followed by a dwell period of 2 minutes, during which the positions of both optical traps were held constant. During the dwell period, the chromosome continued to extend under the applied trapping force, changing the relative position of the bead handles with respect to the centres of the optical traps, which equated to a changing force acting on both beads (Figure 2c). Force acquisition at high frequency (2.5 MHz) was extracted during the dwell period (Figure 2d) and provided continuous broadband viscoelastic data over seven decades of frequencies (10^−2^ −10^5^ rad/s) from single chromosomes.

**Fig. 2.**
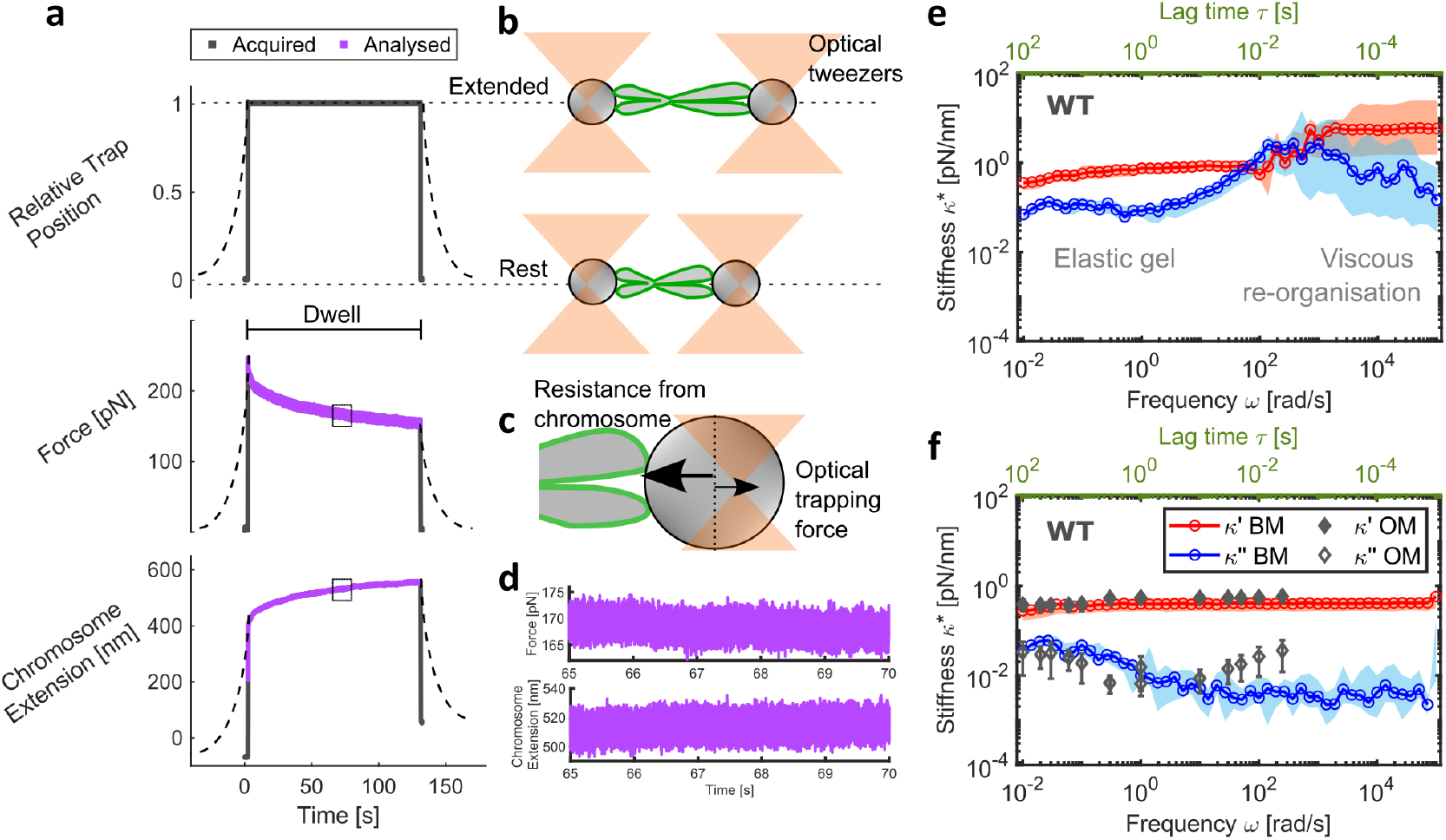
Broadband microrheology of chromosomes. a. Schematic of microrheology experimental procedure. Dashed lines represent trajectory at force-loading rate of 0.2 *µ*m/s and solid lines for 100 *µ*m/s. Data in purple were analysed to provide broadband mechanical response. b. Schematic representation of tweezer and chromosomes positions from (a.) c. Opposing forces experienced by both beads (one shown) in the non-equilibrium state. d. Zoomed-in sub-region of the analysed force and chromosome extension data e. Complex stiffness *κ*^*^(*ω*) with frequency (bottom axis in black) and lag time 1*/ω* (top axis in green) from broadband microrheology (BM) of WT chromosomes at 100 *µ*m/s (median and 95% CI) highlighting regions of viscous reorganisation and gel-like behaviour. Data in blue are the viscous modulus *κ*^*′′*^(*ω*) and in red are the elastic modulus*κ*^*′*^(*ω*) f. *κ*^*^(*ω*) at 0.2 *µ*m/s force-loading rate (median and 95% CI) of WT chromosomes overlaid with oscillatory microrheology (OM) data from Meijering et al., (2022) (27).

This approach far surpasses oscillatory microrheology methods (27, 35) which derive mechanical properties at discrete frequencies only. Our data reveal a fluid-like behaviour at high frequencies (10^1^ − 10^3^ rad/s) following deformation in all tested WT chromosomes (Figure 2e). This fluid-like response likely reflects energy lost to the system from internal friction, probably as a result of the reorganisation of the chromosome network. At lower frequencies, after a characteristic time of approximately 100 ms, the chromosomes equilibrate to a gel-like state dominated by elasticity, with minimal changes in viscoelasticity observed across nearly three decades of frequency.

Measurements were also recorded at a slower force-loading rate of 0.2 *µ*m/s to investigate the influence of loading rate on microrheology on WT chromosomes (Figure 2f). At this reduced loading rate, the reorganisation within the chromosome network occurred faster than the applied strain, resulting in the absence of detectable self-reorganisation dynamics at short timescales. Instead, the measurements revealed a predominantly gel-like response throughout the entire duration of the experiment. Notably, our complex stiffness *κ*^*^(*ω*) values agree well with previously reported oscillatory microrheology measurements of chromosomes (27) over four decades of frequency (Figure 2f). At intermediate frequencies (≃100 rad/s), oscillatory measurements show a similar trend in the viscous modulus *κ*^*′′*^(*ω*) to our ≃100 *µ*m/s measurements. This suggests that the brief cross-over between the *κ*^*′′*^(*ω*) and *κ*^*′*^(*ω*), where viscous self-reorganization processes dominate, might have been captured if oscillatory measurements were continued at higher frequencies than those reported (27) (Figure 2f). These results highlight the need for applying fast force-loading rates and a broadband approach to effectively capture the full spectrum of chromosome molecular biophysics, enabling the study of MCP mechanics in previously unattainable detail.

**Altering MCP levels changes the chromosome viscoelastic response** recorded using our broadband microrheology method with a fast force-loading rate of 100 *µ*m/s (Figure 3a-b and Figure 2e). The ratio of viscous *κ*^*′′*^(*ω*) to elastic *κ*^*′*^(*ω*) components of the chromosome’s mechanical response defines the loss tangent *tanδ*, which provides insight into the force damping or energy dissipation potential of the chromosome. Regardless of MCP status, *tanδ* values for all chromosomes peak at relatively high frequencies (10^2^ − 10^3^ rad/s), where energy dissipation is at maximum due to molecular re-organisation, before reaching a minimum where the chromosome equilibrates to an elastic gel state (Figure 3c). WT chromosomes exhibit a *tanδ* peak value of 2.7 [1.2 5.4; 95%CI]. KD chromosomes show a distinct absence of crossover between *κ*^*′′*^(*ω*) and *κ*^*′*^(*ω*) and consequently, their *tanδ* values remain below 1, peaking at 0.6 [0.4 0.7; 95% CI], indicating a consistent predominantly elastic material with low energy dissipation potential. Higher force damping in the presence of the MCP provides a possible mechanistic explanation for its damage mitigation properties noted previously (22). In contrast, OE chromosomes varied in their response with some exhibiting behaviour similar to WT chromosomes, while others displayed more KD-like behaviour, with overall *tanδ* peak values of 0.5 [0.4 1.6; 95% CI] (Figure 3d). The characteristic time (defined as the inverse of the characteristic frequency) when *tanδ* values peak differs significantly between WT, KD, and OE chromosomes. Most WT chromosomes reach their peak at 10 ms (±1 ms) post-deformation, while KD chromosomes peak earlier at 4 ms (±1 ms). OE chromosomes alternate between these two peak times, with 60% peaking at 4 ms. Additionally, all WT and OE *tanδ* data also show a smaller peak at 4 ms (where not dominant) (Figure 3c), suggesting the presence of two distinct molecular mechanisms driving chromosome relaxation. Consistent with this, the *tanδ* minima for the different conditions were also divergent, with WT chromosomes exhibiting a minimum at 1.5 s (±1 s), KD chromosomes relaxing significantly earlier at 0.2 s (±0.08 s), and OE chromosomes displaying an intermediate relaxation minimum at 0.8 s (±0.5 s). These results collectively indicate MCP-dependent reorganisation mechanisms, which are absent in KD chromosomes lacking the MCP. Furthermore, reorganisation appears to be suppressed in OE chromosomes, suggesting that an optimal MCP load is required for normal mechanical behaviour.

**Fig. 3.**
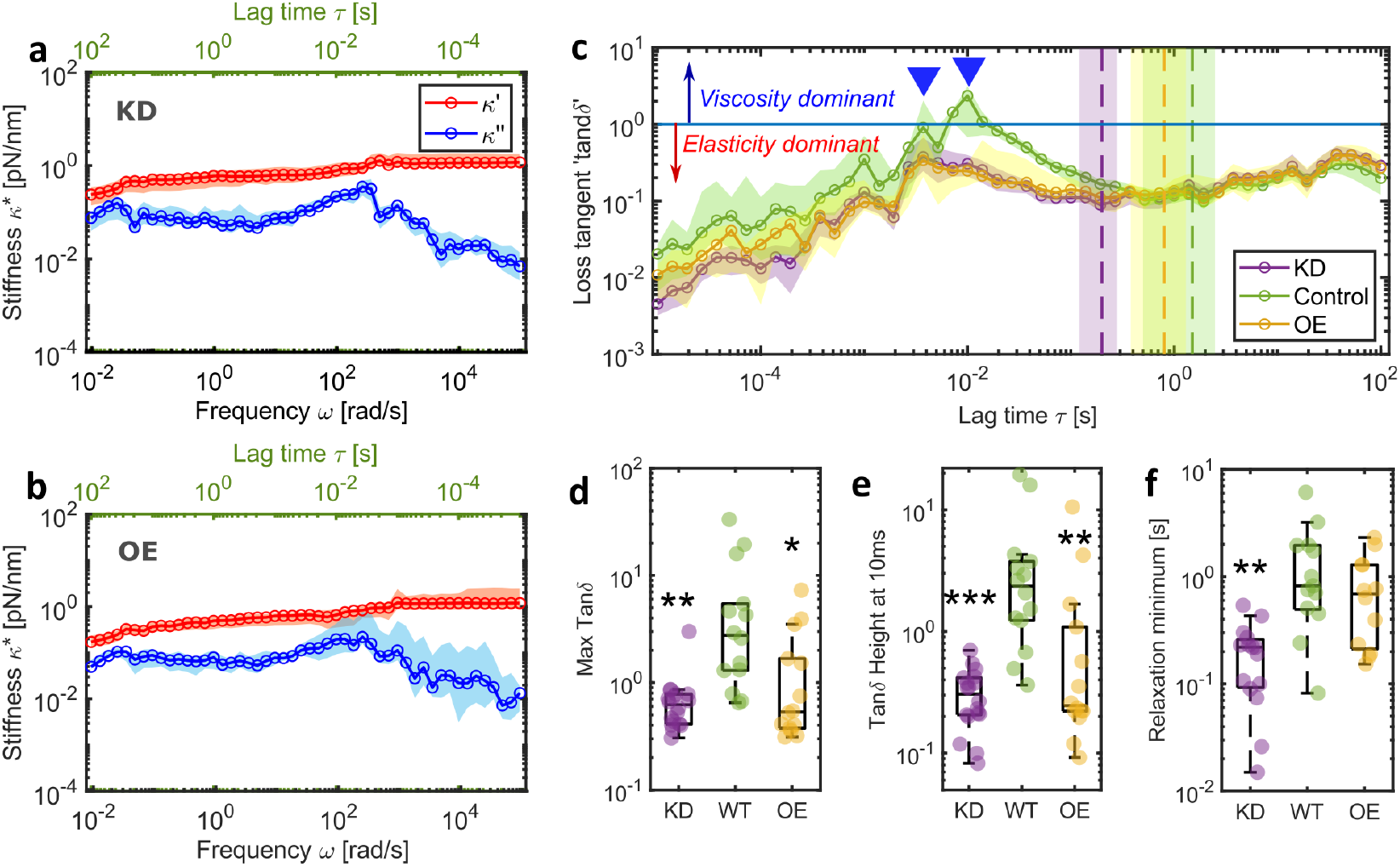
Chromosomes show distinct relaxation dynamics in the presence and absence of the MCP. a. Complex stiffness *κ*^*^(*ω*) of KD chromosomes and b. OE chromosomes compared to WT chromosomes in Figure 2e. c. *tanδ* values for the three conditions (mean ± 95% CI). Blue line represents a *tanδ* value of 1 and values higher than this indicate viscosity is greater than elasticity. Inverted triangles highlight peak reorganisation times with different positions for unaltered and both KD and OE chromosomes. Dashed vertical lines with coloured regions show mean ± 95% CI of time at relaxation minimum. d. Maximum value of *tanδ* for individual chromosomes e. *tanδ* value at 10 ms (position of WT peak) for individual chromosomes and f. *tanδ* minimum after relaxation highlighted by the vertical dashed lines in c. Comparisons to WT (Kruskal-Wallis test), significance values: \**p* < 0.05, \*\**p* < 0.001, \*\*\**p* < 0.0001

The MCP is enriched in proteins with intrinsically disordered domains (14). Ki-67 and NPM1 are notable examples (23, 36) where the disordered domain has been shown to possess patterned charge distributions (36) that enable promiscuous interactions with other proteins and RNA (14, 37). Indeed, unbinding of transient cross-links from electrostatic or van der Waals interactions have been shown to produce pronounced maxima in viscous dissipation at frequencies of 10^−2^ −10^2^ Hz in actin networks (38) and could explain the MCP dependent relaxation dynamics seen in our data. Furthermore, in Ki-67 OE chromosomes with an enriched MCP (Extended Figure S1), an increase in charged domains could alter their viscoelastic properties through electrostatic or steric repulsion to create excluded volumes and reduce molecular flexibility, resulting in gel-like instead of fluid-like behaviour (39). MCP independent reorganisation on the other hand, seen at earlier characteristic times and which appears to be the main relaxation mechanism in KD chromosomes, has been linked to SMC activity (6). The SMC arms in the condensin holocomplex have been reported to be flexible and able to change conformation in millisecond time scales (40), in good agreement with our data (Figure 3c). Other structural proteins such as TopII*α* and Kif4A are also potential candidates involved in this reorganisation and will need to be tested in future studies.

Finally, we sought to fit **a rheological model to parameterize our broadband experimental data** and further isolate the contribution of the MCP to chromosome mechanical behaviour. The Burgers model, previously used to describe the linear viscoelastic properties of nuclei in intact cells (41) and multicellular spheroids (42), was used to interpret chromosome dynamics for the first time (Figure 4a-b). This four-element model consists of two elastic *κ*_1_ and *κ*_2_ and two viscous damping terms *η*_1_ and *η*_2_. *κ*_1_ and *κ*_2_ describe the plateaus in elastic modulus *κ*^*′*^(*ω*) at low and high frequencies respectively, while *η*_1_ in combination with *κ*_1_ and *κ*_2_ characterises the transition in viscoelasticity at intermediate frequencies. *η*_2_ corresponds to the viscosity of the system at long timescales (i.e., low frequencies). Our model fitting indicates that KD chromosomes exhibit reduced *κ*_2_ (Kruskal-Wallis test with multiple comparisons, WT vs KD *p* = 0.002) and *η*_2_ values (Kruskal-Wallis test with multiple comparisons, WT vs KD *p* = 0.02) compared to WT chromosomes, while showing similarities to OE chromosomes. Furthermore, while *κ*_1_ and *η*_1_ remain unchanged regardless of MCP status, the ratio between these two parameters is significantly shifted in KD chromosomes (Kruskal-Wallis test with multiple comparisons, WT vs KD *p* = 0.001), pointing to differences in relaxation processes at intermediate frequencies, probably associated with conformational changes and transient cross-linking. Similarly, the ratio between *η*_2_ and *κ*_1_, relating to viscoelastic behaviour at low frequencies likely driven by large-scale protein interactions, shows significant change in KD chromosomes (Kruskal-Wallis test with multiple comparisons, WT vs KD *p* = 0.02). The elastic parameters extracted from the model align with our single-frequency force-extension experiments, which reveal a lower relative change in stiffness in KD chromosomes (Figure 4e and 1e; Kruskal-Wallis with multiple comparisons of relative *κ* from model fitting, WT vs KD *p* = 0.0004), but no statistically significant differences with MCP status in elasticity at low frequencies (Figure 4c and 1d). The Burgers model can be interpreted as a combination of a Kelvin-Voigt solid, expressed by *κ*_1_ and *η*_1_, likely reflecting the contribution of the chromosome as a whole, and a Maxwell liquid, represented by *κ*_2_ (instantaneous response) and *η*_2_ (long time viscous flow), which may capture the contribution of the MCP coat among other chromosomal elements (Figure 4f). Notably, although Ki-67 and the MCP have often been described as exhibiting ‘liquid-like’ properties (18, 23, 36), our data represents the first direct measurements of these properties.

**Fig. 4.**
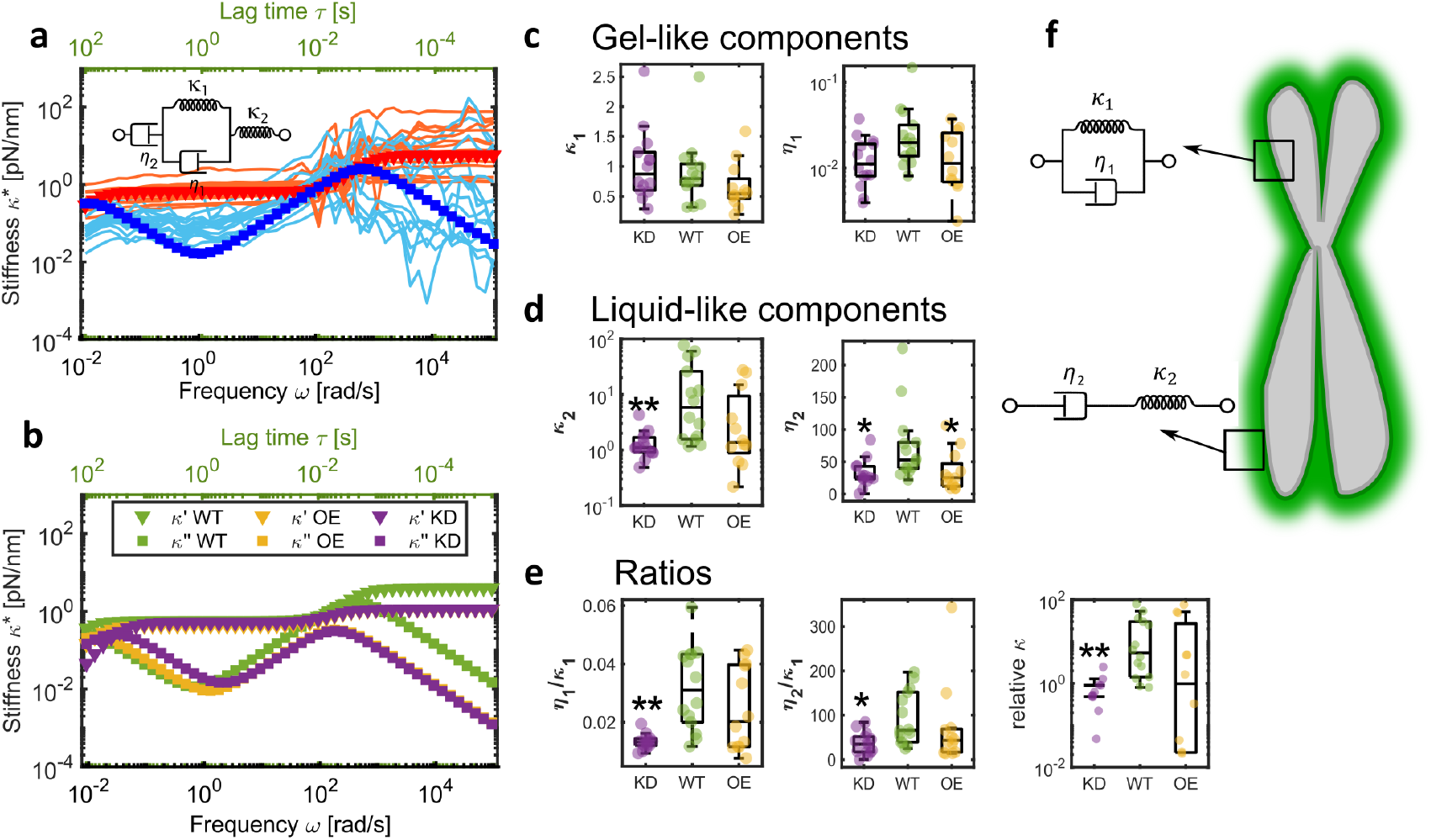
Chromosome mechanical behaviour explained using the Burgers model. a. Burgers model fit to experimental data of individual WT chromosomes with inset four-element spring-dashpot diagram. b. Model fits to average WT, KD and OE data. c-d Comparisons of parameters extracted by fitting experimental data to the Burgers model individually. e. Ratios of fit parameters. f, Schematic of a chromosome to show gel-like properties are associated with the whole chromosome while the MCP behaves like a liquid. For c, d and e) comparisons to WT (Kruskal-Wallis test), significance values: \**p* < 0.05, \*\**p* < 0.001

The chromosome periphery has been shown to be self-supporting, retaining its structure even after DNA and RNA digestion (43). While removing the MCP by silencing Ki-67 expression does not prevent chromosome condensation, its disruption at a later stage of mitosis results in distorted chromosomes (20). The precise nature of linkages that maintain the MCP compartment remains unclear, however, it has been shown to function independently of TopoII*α* and condensins (20). Our data provide the first direct evidence of the influence of the MCP on stiffening behaviour of individual chromosomes, as further support for a periphery-based structural constraint on the chromosome. Transient cross-links driven by charged interactions with intrinsically disordered proteins like Ki-67 may explain the dynamic, ‘fluid’ nature of this network. Such interactions have recently been shown to influence chromosome fluidity by altering contour length (35). However, a Ki-67-driven structural constraint has previously been dismissed in favour of nucleosome-nucleosome interactions at the chromosome periphery (44). These nucleosome interactions, however, do not fully account for our findings, which highlight a distinct MCP-driven chromosome relaxation process, warranting further investigation.

In summary, this study highlights the importance of broadband analysis for capturing the time-dependent dynamics of chromosomes. Our data reveal a timeline of dynamic mechanical events following chromosome stretching, spanning timescales from tens of microseconds to minutes, and elucidate the direct role of the MCP in regulating chromosome fluidity and structural integrity. Increased Ki-67 expression is a hallmark of cancer progression and there is strong interest in using this protein as a therapeutic target (45–47) paving a translational path for our findings. Furthermore, temporal mapping of mechanical responses provides a foundation for exploring the role of charged interactions in chromosomes and more broadly, the biophysics of phase separation, another emerging therapeutic target (48). Finally, broadband rheology of single chromosomes opens avenues for investigating the contributions of other chromosome associated structures, many of which are still poorly defined.

## Methods

### Generation of a HeLa Ki-67-mEmerald cell line

DNA oligonucleotides (Forward 5’ ACATGGACATGAGCCCCCTG 3’, Reverse 5’ GATAGTTCTGGGGCCTCAGG 3’) used for gRNA synthesis were designed using the Benchling CRISPR gRNA Design Tool available at (www.benchling.com, accessed on 15 May 2022) and ordered through Sigma. gRNA was annealed together and cloned into the Bbs1 restriction site of the `pX330-U6-Chimeric_BB-CBh-hSpCAas9-hGem` (1/110) vector (Addgene #71707) before transformation into NEB® 5-alpha Competent E. coli (New England Biolabs, Ipswich, Massachusetts, USA). DNA was prepared from single colonies before sequence verification using an Applied Biosystems™ (Waltham, Massachusetts, USA) 3130xl genetic analyser. Homology-directed repair (HDR) plasmids designed to introduce mEmerald to the C-terminus of MKI67 were synthesised using a pUC18 backbone and ordered from GenScript (Piscataway, NJ, USA). The plasmids also included a puromycin resistance cassette to facilitate efficient positive selection of transfected cells, as well as a T2A sequence to separate the puromycin-resistant protein from the mEmerald-fused Ki-67 protein. HeLa cells were nucleofected with the HDR plasmid and the gRNA cas9 plasmid and allowed to recover for 3 days before selection with G418 for 7 days. Cells were then treated with puromycin (Fisher Scientific) for 3 days and single cell sorted into ninety-six well plates. Clones were screened by PCR genotyping.

### Overexpression Construct

A gRNA sequence targeting the MKI67 gene was cloned into a CRISPRa plasmid (Addgene #175572) containing an inactivated CRISPR/Cas9 fusion protein with the VPR transcriptional activation domain, and mCherry as a fluorescent reporter.

### Tissue culture and transfection

The Ki-67-mEmerald cell line was maintained in DMEM supplemented with 10% fetal bovine serum (FBS) and 1% antibiotics (penicillin-streptomycin) at 37°C in 5% CO2 and was regularly tested for mycoplasma contamination. To knock out Ki-67, the Ki-67-mEmerald cell line was transiently transfected with 0.8 *µ*g siRNA (Ki-5 in 13) per mL of growth media using Lipofectamine RNAiMax (Invitrogen) according to manufacturer’s instructions and analysed after 72 hours. Ki-67 was transiently overexpressed in the Ki-67-mEmerald cell line with 0.8 *µ*g of the overexpression construct per mL of growth media using jetPRIME (VWR, Lutterworth, Leicestershire, UK) transfection reagent (PolyPlus).

### Chromosome isolation

Ki-67-mEmerald cells were grown to exponential growth phase in T125 flasks before transfection for 72 hours to knock out or overexpress Ki-67. Growth media was supplemented with biotin (50 mM) 24 hours before chromosome isolation using a protocol similar to previously described (27). Briefly, cells were arrested with Nocadazole (100 ng/mL) treatment for 14 hours to enrich for mitosis. Arrested cells were collected by mitotic shake-off, centrifuged at 300*g* for 5 min and incubated in a swelling buffer (10 mL of 75 mM KCl and 5 mM Tris-HCl, pH 8.0 for 107 cells) for 30 min at 37 °C. The swollen cells were then centrifuged at 4 °C and re-suspended in 8 mL of cold polyamine (PA) buffer (15 mM Tris-HCl (pH 8.0), 2 mM EDTA, 0.5 mM EGTA, 80 mM KCl, 20 mM NaCl, 0.5 mM spermidine, 0.2 mM spermine and 0.2% Tween-20) before mechanically breaking in the presence of protease inhibitors (Pierce) and phosphatase inhibitors (PhosSTOP, Roche) in a 15 mL dounce homogeniser (Kimble) with 40 strokes of a tight pestle on ice. The homogenised suspension was cleared of debris three times before glycerol gradient fractionation (2 mL each of 60% and 30% glycerol in PA) by centrifugation at 1750*g* for 30 min at 4 °C. Isolated chromosomes were collected from the 60% glycerol fraction. Chromosomes were stored in 60% glycerol at -20 °C and used within three months of isolation.

### Chromosome micromanipulations

A C-trap Edge-450 (LUMICKS) instrument at the University of Nottingham and another at the University of Kent were used to analyse chromosome dynamics. This instrument combines optical tweezers (up to 4 optical traps) and microfluidics with multichannel laminar flow which allowed for the incremental assembly of the experimental unit (Extended Figure S2). The instrument was used in the dual-trap mode with a 50% split of laser power between the two optical traps. A pair of streptavidin coated polystyrene beads (3 *µ*m diameter, Bangs Laboratories) were optically trapped in the first microfluidic channel with the two optical traps. Trap stiffness was calibrated for every new bead pair in the absence of fluid flow and was 0.5 ± 0.03 pN/nm at each bead. The trapped bead pair was then moved across to the channel flowing chromosomes suspended in PA buffer (channel 2) to capture a single biotinylated chromosome first by binding to one bead. The fluid flow oriented the attached chromosome such that a free end was available for the second bead to be brought into contact with it to form a dumbbell unit. Chromosome attachment was biased along the telometric ends due to the orientation of the chromosomes within the fluid flow. Non-biotinylated chromosomes did not attach to the beads. Experimental manipulations were performed using this dumbbell configuration aligned along the X-axis of the C-trap imaging system, in a third microfluidic channel containing only PA buffer as detailed in the sections below.

Fluorescence images were simultaneously acquired during experiments using HILO (pseudo TIRF) microscopy on the C-trap. Intensities of individual chromosomes (see image analysis section) were used to exclude chromosomes from untransfected cells within each treatment group. Only chromosomes between 2-5 *µ*m in length were analysed to ensure single chromosomes with telomeric attachment were being tested. Analysis was performed in MATLAB 2022b (MathWorks) using custom scripts.

### Force-extension experiments

Single frequency force-extension experiments (Figure 1) involved keeping one optical trap in the dumbbell unit stationary while linearly displacing the second trap position along the X-axis at a fixed rate of 0.02 *µ*m/s to measure linear elasticity and 0.2 *µ*m/s to examine chromosome non-linear stiffening behaviour using the inbuilt force spectroscopy module in Bluelake (LUMICKS) which removed subjectivity during chromosome manipulation. Chromosome response to stretching is variable (Figure 1c) so only chromosomes showing linear behaviour at 0.02 *µ*m/s and non-linear behaviour at 0.2 *µ*m/s were analysed. In each case, the optical trap was continuously displaced until the resistance from the chromosome was greater than that could be overcome by the optical trapping force, resulting in one or both beads leaving the optical trap.

The stretch modulus *S* was computed as the slope of force against normalised chromosome extension, (*L*(*t*) − *L*_0_)*/L*_0_ , where *L*(*t*) is the length of the chromosome at the given time and *L*_0_ is its initial untangled length before any extension was registered as an increase in force. Non-linear stiffening analysis was performed as described previously (27). In brief, data were smoothed using a moving average function with a window size 1/15 of the data points in the measurement. Stiffening exponent *γ* was computed by fitting an exponential function of the form

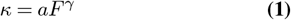

to the numerical gradient *κ* of the force-extension curve against force *F*.

### Microrheology

A new broadband microrheology technique was developed to analyse the time-dependent response of single chromosomes to a stretching force (Figure 2). At the start of the experiment, the chromosome was fully extended in the chromosome dumbbell configuration such that any further extension by moving one of the bead handles registered an increase in force acting on each bead. Force was exerted on the captured chromosome by laterally displacing the position of one of the two optical traps by 1 *µ*m at speeds of 100 *µ*m/s. To examine the effect of force-loading rates on chromosome response, force-loading at 0.2 *µ*m/s was also performed. In these experiments, the moving optical trap was displaced until a force readout of 150 pN on the bead was detected. The chromosome was then held under stress in the extended position for around 2 min before restoring it back to its original state by returning the displaced optical trap to the start position. High frequency (2.5 MHz) force data over the 2 min dwell period was used in the microrheology analysis. Data collection was automated using the force distance module in Bluelake (LUMICKS).

The force acting at each optically trapped bead can be calculated as

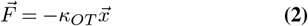

where 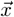 is the displacement of the bead from the optical trap centre and *κ*_*OT*_ is the stiffness of the optical trap.

The net stretching force *F*(*t*) is the cumulative force acting on the chromosome and is measured as the absolute sum of the force acting at both beads. The strain *ϵ*(*t*) is calculated as the relative extension of the chromosome:

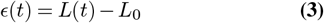

The high frequency position information for each optically trapped bead was computed by subtracting 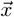 from the position of the optical trap centre holding that bead.

The broadband mechanical response of individual chromosomes was computed as complex stiffness

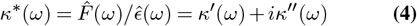

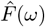 and 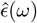 are the Fourier transforms of *F*(*t*) and *ϵ*(*t*) respectively.

The loss tangent *tanδ* (Figure 3c) was calculated as

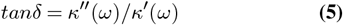

*κ*^*^(*ω*) and *tanδ* were acquired using a MATLAB application i-Rheo C-Stretch (see Code availability and Extended Figure S3) that implements a Fourier transform method introduced previously (49, 50). The characteristic minimum time in *tanδ* was extracted as the time at which an exponential fit to the data from each individual chromosome reached the minimum value.

### Model fitting

The Kelvin representation of the Burgers model can be expressed as,

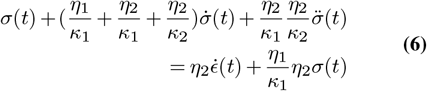

Here, *κ*_1_, *κ*_2_, *η*_1_ and *η*_2_ are the elastic and viscous parameters in the model respectively and can be diagrammatically represented by the spring and dashpot schematic in Figure 4a. Solving the Burgers model for complex stiffness from equations (4) and (6) gives,

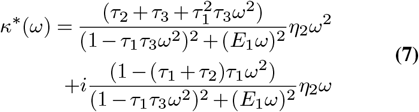

where, *τ*_1_ = *η*_1_*/κ*_1_ , *τ*_2_ = *η*_2_*/κ*_1_ , *τ*_3_ = *η*_2_*/κ*_2_ and *E*_1_ = *τ*_1_ + *τ*_2_ +*τ*_3_

Individual microrheology data from experiments were fit to the above equation using non-linear least squared fitting in MATLAB to extract *κ*_1_, *κ*_2_, *η*_1_ and *η*_2_ (Figure 4).

### Indirect immunofluorescence and microscopy

Primary antibodies were used as follows: NPM1 (Mouse monoclonal, #32-5200 (Thermo fisher, Waltham, Massachusetts, USA) 1:60; Nucleolin (Rabbit polyclonal, Abcam, Cambridge, UK, #ab22758) 1:100. Fluorescence-labelled secondary antibodies were applied at 1:400 (Jackson ImmunoResearch, Ely, UK). For immunofluorescence, cells were fixed in 3.5% paraformaldehyde for 15 min, permeabilized in 0.3% Triton X-100 for 5 min and blocked in 2% blocking solution (Thermo fisher). Cells were incubated overnight with the primary antibodies washed in PBS and secondary antibodies were applied for 1 hr before counter-staining with DAPI. Fluorescent image acquisition was performed using a Leica (Wetzlar, Germany) TCS SPE confocal microscope with an x63 oil immersion objective. Microscope settings including laser power, exposure time and pinhole size were kept constant across all images in Extended Figure S1. Images were exported as TIFF files.

### Image Analysis

Quantification of fluorescence intensity was performed in ImageJ (51). For chromosome images from single-chromosome experiments (Figure 1b), fluorescence intensity was measured as the integrated density from line scans across each chromosome, divided by the length of the chromosome. NPM1 and nucleolin levels were quantified with Ki-67 manipulation from multichannel immunofluorescence microscopy images of early mitotic (prometaphase and metaphase) cells. Masks were generated by thresholding the DAPI channel to outline chromosomes and the NPM1 or nucleolin channel respectively to mark the spread of these proteins within the cell. The perichromosomal levels of NPM1 and nucleolin were measured as the mean intensity within the intersection of the two masks (using the AND function on the masks in the Image Calculator settings) in the respective image channel. Ki-67 levels were measured as the mean intensity within the chromosome mask in the mEmerald channel.

### Calculations and Statistics

Kruskal-Wallis tests with post-analysis multiple comparisons and linear regression were performed in MATLAB (2022b, MathWorks). *P* values of ≤ 0.05 were considered to represent significance.

## Code availability

The i-Rheo C-Stretch app can be downloaded from https://github.com/tvmendonca/iRheoCStretch (Extended Figure S3)

## Data availability

Data will be made available at the time of peer-reviewed publication.

## ACKNOWLEDGEMENTS

This work was funded by a Leverhulme Trust research project grant (RPG-2021-118) awarded to DGB and AJW. Data acquisition at the University of Kent was enabled by a Travelling Fellowship to TM from the Company of Biologists (JC-STF23081207). The LUMICKS C-trap instrument at University of Kent was funded by BBSRC (`BB/T017767/1`) to NMK and one at the University of Nottingham was funded by a BBSRC Alert grant (`BB/X019837/1`) to DGB, TM and AJW. DGB thanks; BBSRC David Phillips Fellowship (`BB/V005626/1`), Royal Society (`RGS/R2/202366`), Academy of Medical Sciences Springboard Award (`SBF006\1071`), and the Wellcome Trust (GC, Drug Discovery and Team Science DTP `218466/Z/19/Z`).

## Extended Data

**Fig. S1.**
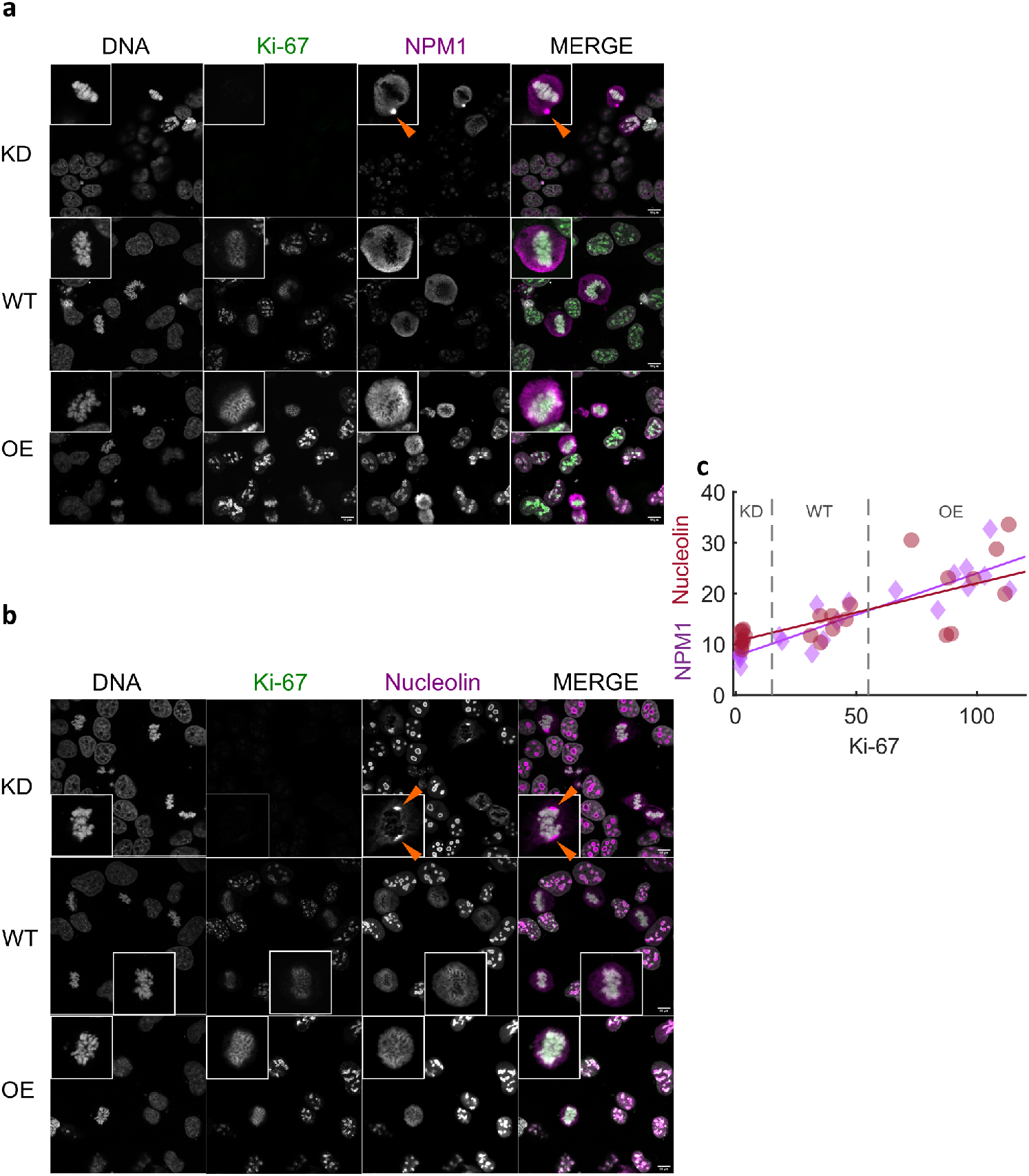
Ki-67 levels dictate localisation of the periphery proteins nucleophosmin (NPM1) and nucleolin at mitotic chromosomes. Confocal images of Ki-67-mEmerald cells with wild type (WT), knockdown (KD) or overexpressed (OE) levels of Ki-67, immuno-labelled for a. NPM1 and b. nucleolin. Cells were counterstained for DNA with DAPI. NPM1 and nucleolin aggregates (orange arrows) in Ki-67 KD mitotic cells. Insets show zoomed in individual mitotic cells from images. Scale bar = 10 *µ*m. c. Mean fluorescence intensity (arbitrary units) of NPM1 and nucleolin at the chromosome periphery vs intensity of Ki-67 in early mitotic WT, KD and OE Ki-67-mEmerald cells with linear regression fits (NPM1 vs Ki-67: *r*^2^ = 0.8, *p* < 0.0001 and nucleolin vs Ki-67: *r*^2^ = 0.6, *p* < 0.0001)

**Fig. S2.**
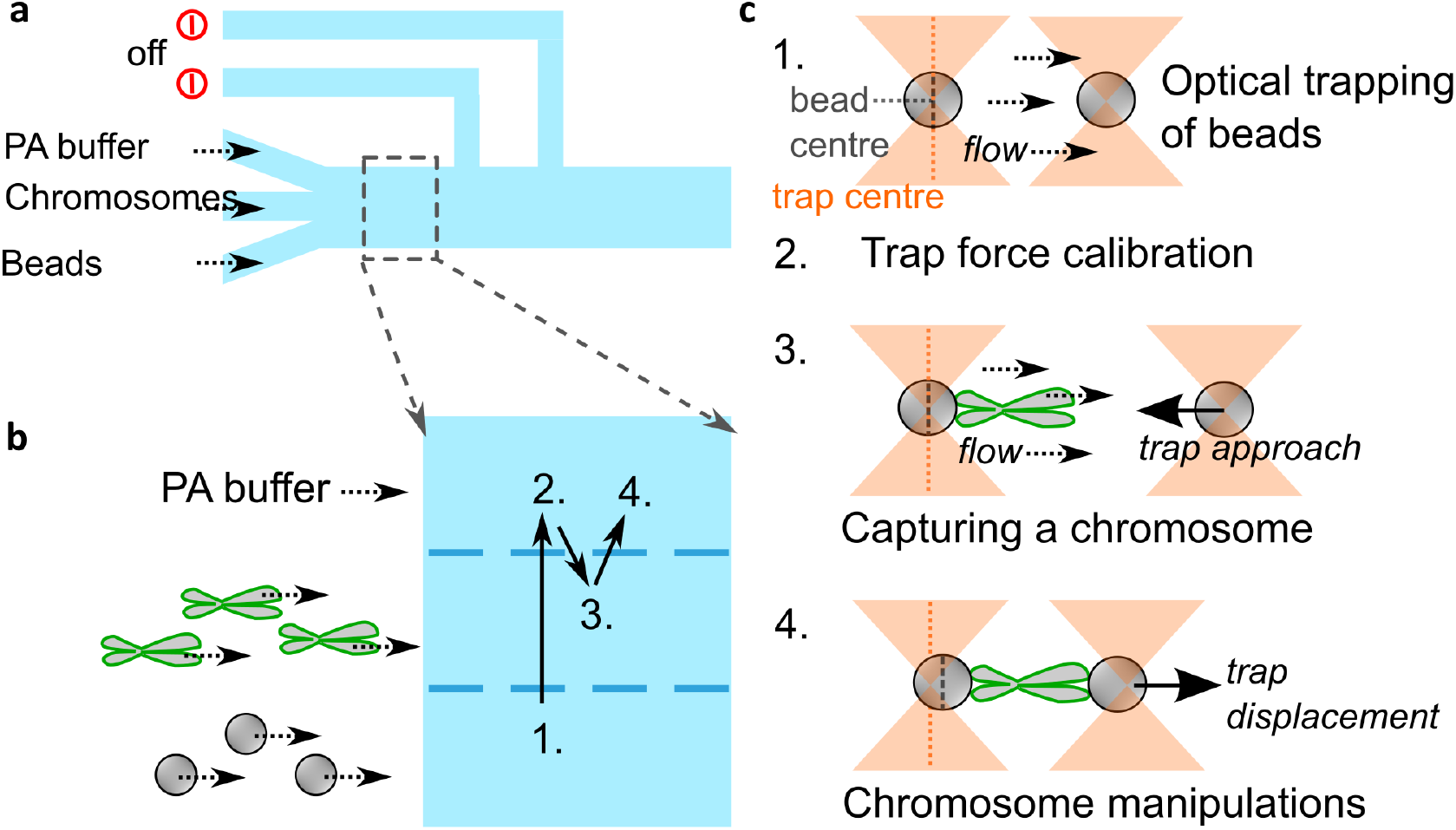
Assembling the chromosome dumbbell unit for single-chromosome manipulations. a. Schematic of the u-Flux microfluidics flow cell (type c1). Only three channels were used. b. Sub-region of the flow cell mapping the positions of each step in dumbbell assembly procedure. c. The four main steps to all chromosome manipulation experiments: 1. Optical trapping of beads as they flow through channel 1. Dotted vertical lines represent centres of the optical trap (orange) and bead (grey), only shown on one side but applies to both. Dotted arrows represent direction of fluid flow. 2. Each new pair of trapped beads were calibrated to accurately measure trapping force on both beads. Fluid flow is switched off during calibration. 3. A single chromosome is captured by binding to one bead first and then bringing the second bead into contact to form the dumbbell. 4. Single chromosomes are stretched in the absence of fluid flow by displacing the position of one optical trap while leaving the second stationary. The stretched chromosome displaces the bead handles such that the bead centres and optical trap centres are no longer coincident.

**Fig. S3.**
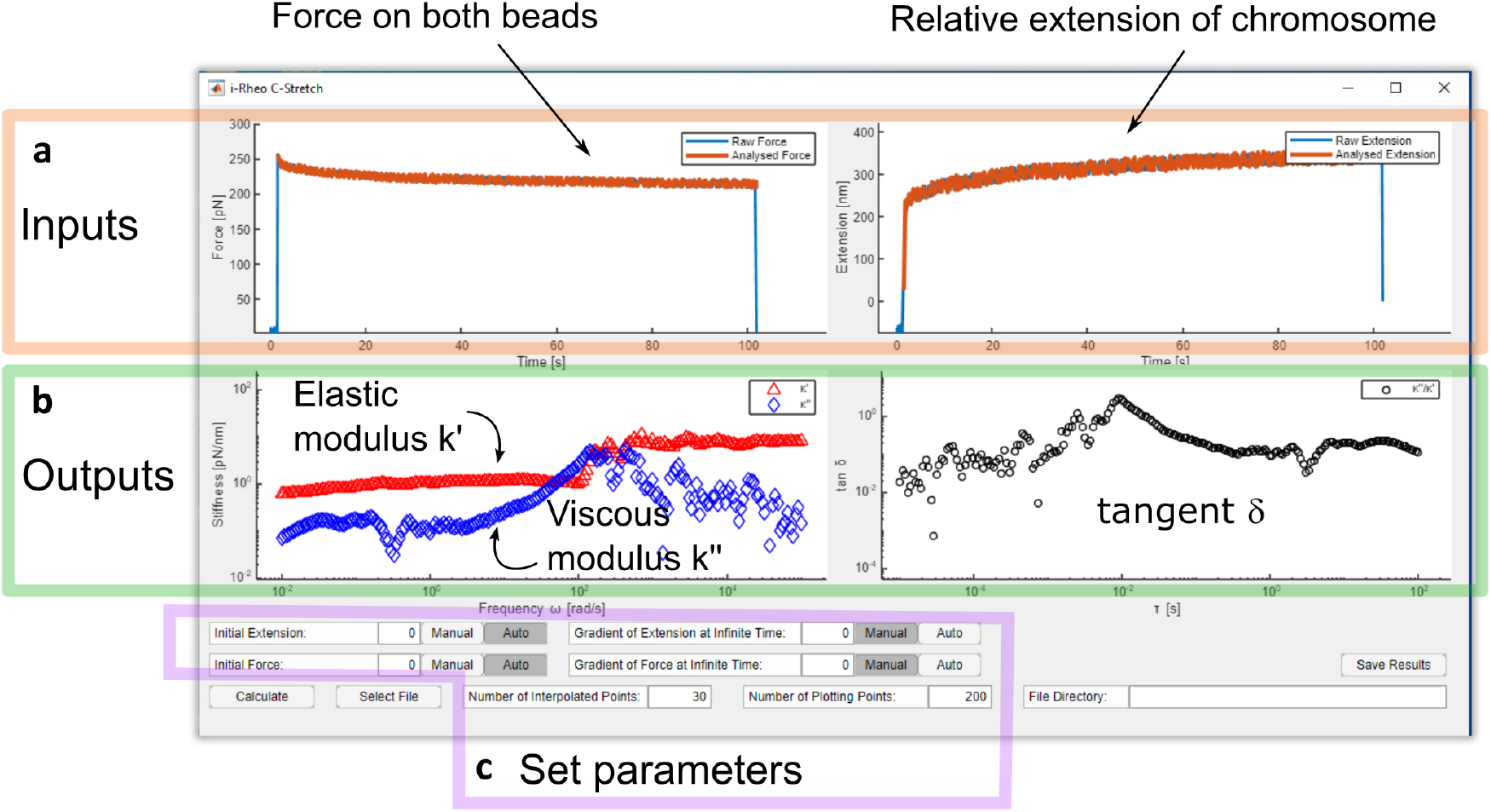
Screenshot of the i-Rheo C-Stretch app. This plug and play app accepts 3xn datasets of time, summed force on both beads and relative extension of chromosome. a. The inputs are displayed in the two plots at the top and b. the output complex stiffness and *tanδ* results are displayed in the plots at the bottom of the app. c. Users can toggle between auto detection of parameters (initial and gradient values for force and extension) or manual setting if known along with adjusting the number of interpolation points and plotting density. Complex stiffness values can be exported from the app. A detailed guide is available to download with the app (see code availability statement)

## Bibliography

1. M.G. Poirier and J.F. Marko. Micromechanical studies of mitotic chromosomes. 23(5):409– 431. ISSN 1573-2657. doi: 10.1023/A:1023402321367.

2. T. Man, H. Witt, E. J. G. Peterman, and G. J. L. Wuite. The mechanics of mitotic chromosomes. 54:e10. ISSN 0033-5835, 1469-8994. doi: 10.1017/S0033583521000081.

3. Veer I. P. Keizer, Simon Grosse-Holz, Maxime Woringer, Laura Zambon, Koceila Aizel, Maud Bongaerts, Fanny Delille, Lorena Kolar-Znika, Vittore F. Scolari, Sebastian Hoffmann, Edward J. Banigan, Leonid A. Mirny, Maxime Dahan, Daniele Fachinetti, and Antoine Coulon. Live-cell micromanipulation of a genomic locus reveals interphase chromatin mechanics. 377(6605):489–495. doi: 10.1126/science.abi9810. Publisher: American Association for the Advancement of Science.

4. Manon Valet and Angelo Rosa. Viscoelasticity of model interphase chromosomes. 141(24):245101. ISSN 0021-9606. doi: 10.1063/1.4903996.

5. S. Almagro, S. Dimitrov, T. Hirano, M. Vallade, and D. Riveline. Individual chromosomes as viscoelastic copolymers. 63(6):908, . ISSN 0295-5075. doi: 10.1209/epl/i2003-00609-3. Publisher: IOP Publishing.

6. Benjamin S. Ruben, Sumitabha Brahmachari, Vinícius G. Contessoto, Ryan R. Cheng, Antonio B. Oliveira Junior, Michele Di Pierro, and José N. Onuchic. Structural reorganization and relaxation dynamics of axially stressed chromosomes. 122(9):1633–1645. ISSN 0006-3495. doi: 10.1016/j.bpj.2023.03.029.

7. John F. Marko. Micromechanical studies of mitotic chromosomes. 16(3):469. ISSN 1573-6849. doi: 10.1007/s10577-008-1233-7.

8. Sébastien Almagro, Daniel Riveline, Tatsuya Hirano, Bahram Houchmandzadeh, and Stefan Dimitrov. The mitotic chromosome is an assembly of rigid elastic axes organized by structural maintenance of chromosomes (SMC) proteins and surrounded by a soft chromatin envelope*. 279(7):5118–5126, . ISSN 0021-9258. doi: 10.1074/jbc.M307221200.

9. Mingxuan Sun, Ronald Biggs, Jessica Hornick, and John F. Marko. Condensin controls mitotic chromosome stiffness and stability without forming a structurally contiguous scaffold. 26(4):277–295. ISSN 1573-6849. doi: 10.1007/s10577-018-9584-1.

10. Daniel G. Booth and William C. Earnshaw. Ki-67 and the chromosome periphery compartment in mitosis. 27(12):906–916. ISSN 0962-8924. doi: 10.1016/j.tcb.2017.08.001.

11. Danièle Hernandez-Verdun and Thierry Gautier. The chromosome periphery during mitosis. 16(3):179–185. ISSN 1521-1878. doi: 10.1002/bies.950160308. _eprint: https://onlinelibrary.wiley.com/doi/pdf/10.1002/bies.950160308.

12. Chas. W. Metz and José F. Nonidez. The behavior of the nucleus and chromosomes during spermatogenesis in the robber fly lasiopogon bivittatus. 46(4):153–164. ISSN 0006-3185. doi: 10.2307/1536505. Publisher: Marine Biological Laboratory.

13. Daniel G Booth, Masatoshi Takagi, Luis Sanchez-Pulido, Elizabeth Petfalski, Giulia Vargiu, Kumiko Samejima, Naoko Imamoto, Chris P Ponting, David Tollervey, William C Earnshaw, and Paola Vagnarelli. Ki-67 is a PP1-interacting protein that organises the mitotic chromosome periphery. 3:e01641. ISSN 2050-084X. doi: 10.7554/eLife.01641. Publisher: eLife Sciences Publications, Ltd.

14. Lovisa Stenström, Diana Mahdessian, Christian Gnann, Anthony J. Cesnik, Wei Ouyang, Manuel D. Leonetti, Mathias Uhlén, Sara Cuylen-Haering, Peter J. Thul, and Emma Lundberg. Mapping the nucleolar proteome reveals a spatiotemporal organization related to intrinsic protein disorder. 16(8):e9469. ISSN 1744-4292. doi: 10.15252/msb.20209469.

15. Emily Sparago, Reito Watanabe, Judith A. Sharp, and Michael D. Blower. Dynamic redistribution and inheritance of chromatin:RNA interactions during cell division. 1. ISSN 2813-7116. doi: 10.3389/frnar.2023.1240954. Publisher: Frontiers.

16. Sara Cuylen, Claudia Blaukopf, Antonio Z. Politi, Thomas Müller-Reichert, Beate Neumann, Ina Poser, Jan Ellenberg, Anthony A. Hyman, and Daniel W. Gerlich. Ki-67 acts as a biological surfactant to disperse mitotic chromosomes. 535(7611):308–312. ISSN 1476-4687. doi: 10.1038/nature18610. Bandiera_abtest: a Cg_type: Nature Research Journals Number: 7611 Primary_atype: Research Publisher: Nature Publishing Group Subject_term: Biopolymers in vivo;Cancer;Chromosome segregation;Chromosomes Subject_term_id: biopolymers-in-vivo;cancer;chromosome-segregation;chromosomes.

17. Kumiko Samejima, Daniel G. Booth, Hiromi Ogawa, James R. Paulson, Linfeng Xie, Cara A. Watson, Melpomeni Platani, Masato T. Kanemaki, and William C. Earnshaw. Functional analysis after rapid degradation of condensins and 3d-EM reveals chromatin volume is uncoupled from chromosome architecture in mitosis. 131(4). ISSN 0021-9533. doi: 10.1242/jcs.210187.

18. Alberto Hernandez-Armendariz, Valerio Sorichetti, Yuki Hayashi, Zuzana Koskova, Andreas Brunner, Jan Ellenberg, Anđ ela Šarić, and Sara Cuylen-Haering. A liquid-like coat mediates chromosome clustering during mitotic exit. 84(17):3254–3270.e9. ISSN 1097-2765. doi: 10.1016/j.molcel.2024.07.022. Publisher: Elsevier.

19. Konstantinos Stamatiou and Paola Vagnarelli. Chromosome clustering in mitosis by the nuclear protein ki-67. 49(6):2767–2776. ISSN 0300-5127. doi: 10.1042/BST20210717.

20. Masatoshi Takagi, Takao Ono, Toyoaki Natsume, Chiyomi Sakamoto, Mitsuyoshi Nakao, Noriko Saitoh, Masato T. Kanemaki, Tatsuya Hirano, and Naoko Imamoto. Ki-67 and condensins support the integrity of mitotic chromosomes through distinct mechanisms. 131(6):jcs212092. ISSN 0021-9533. doi: 10.1242/jcs.212092.

21. Konstantinos Stamatiou, Florentin Huguet, Lukas V. Serapinas, Christos Spanos, Juri Rappsilber, and Paola Vagnarelli. Ki-67 is necessary during DNA replication for fork protection and genome stability. 25:105. doi: 10.1186/s13059-024-03243-5.

22. Osama Garwain, Xiaoming Sun, Divya Ramalingam Iyer, Rui Li, Lihua Julie Zhu, and Paul D. Kaufman. The chromatin-binding domain of ki-67 together with p53 protects human chromosomes from mitotic damage. 118(32). ISSN 0027-8424, 1091-6490. doi: 10.1073/pnas.2021998118. ISBN: 9782021998115 Publisher: National Academy of Sciences Section: Biological Sciences.

23. Hiroya Yamazaki, Masatoshi Takagi, Hidetaka Kosako, Tatsuya Hirano, and Shige H. Yoshimura. Cell cycle-specific phase separation regulated by protein charge blockiness. 24(5):625–632. ISSN 1476-4679. doi: 10.1038/s41556-022-00903-1. Number: 5 Publisher: Nature Publishing Group.

24. A. Ashkin, J. M. Dziedzic, J. E. Bjorkholm, and Steven Chu. Observation of a single-beam gradient force optical trap for dielectric particles. 11(5):288. ISSN 0146-9592. doi: 10.1364/OL.11.000288. Publisher: Optical Society of America ISBN: 9781482269017.

25. Michael G. Poirier, Ajay Nemani, Prateek Gupta, Sertac Eroglu, and John F. Marko. Probing chromosome structure with dynamic force relaxation. 86(2):360–363, . ISSN 0031-9007, 1079-7114. doi: 10.1103/PhysRevLett.86.360.

26. Michael G. Poirier, Sertac Eroglu, and John F. Marko. The bending rigidity of mitotic chromosomes. 13(6):2170–2179, . ISSN 1059-1524.

27. Anna E. C. Meijering, Kata Sarlós, Christian F. Nielsen, Hannes Witt, Janni Harju, Emma Kerklingh, Guus H. Haasnoot, Anna H. Bizard, Iddo Heller, Chase P. Broedersz, Ying Liu, Erwin J. G. Peterman, Ian D. Hickson, and Gijs J. L. Wuite. Nonlinear mechanics of human mitotic chromosomes. 605(7910):545–550. ISSN 1476-4687. doi: 10.1038/s41586-022-04666-5.

28. Goran Žagar, Patrick R. Onck, and Erik van der Giessen. Two fundamental mechanisms govern the stiffening of cross-linked networks. 108(6):1470–1479. ISSN 0006-3495. doi: 10.1016/j.bpj.2015.02.015. Publisher: Elsevier.

29. John F. Marko and Eric D. Siggia. Stretching DNA. 28(26):8759–8770. ISSN 0024-9297. doi: 10.1021/ma00130a008. Publisher: American Chemical Society.

30. Masao Doi, Sam F. Edwards, and Samuel Frederick Edwards. The Theory of Polymer Dynamics. Clarendon Press. ISBN 978-0-19-852033-7. Google-Books-ID: dMzGyWs3GKcC.

31. Michael Poirier, Sertac Eroglu, Didier Chatenay, and John F. Marko. Reversible and irreversible unfolding of mitotic newt chromosomes by applied force. 11(1):269–276, . ISSN 1059-1524. doi: 10.1091/mbc.11.1.269. Publisher: American Society for Cell Biology(mboc).

32. Feroz M. Hameed, Madan Rao, and G. V. Shivashankar. Dynamics of passive and active particles in the cell nucleus. 7(10):e45843. ISSN 1932-6203. doi: 10.1371/journal.pone.0045843. Publisher: Public Library of Science.

33. Yiider Tseng, Jerry S. H. Lee, Thomas P. Kole, Ingjye Jiang, and Denis Wirtz. Microorganization and visco-elasticity of the interphase nucleus revealed by particle nanotracking. 117(10):2159–2167. ISSN 0021-9533. doi: 10.1242/jcs.01073.

34. Ya Hua Chim, Louise M. Mason, Nicola Rath, Michael F. Olson, Manlio Tassieri, and Huabing Yin. A one-step procedure to probe the viscoelastic properties of cells by atomic force microscopy. 8. doi: 10.1038/s41598-018-32704-8. Publisher: Nature Publishing Group.

35. Hannes Witt, Janni Harju, Emma M. J. Chameau, Charlotte M. A. Bruinsma, Tinka V. M. Clement, Christian F. Nielsen, Ian D. Hickson, Erwin J. G. Peterman, Chase P. Broedersz, and Gijs J. L. Wuite. Ion-mediated condensation controls the mechanics of mitotic chromosomes. 23(11):1556–1562. ISSN 1476-4660. doi: 10.1038/s41563-024-01975-0. Publisher: Nature Publishing Group.

36. Lucy Remnant, Natalia Y. Kochanova, Caitlin Reid, Fernanda Cisneros-Soberanis, and William C. Earnshaw. The intrinsically disorderly story of ki-67. 11(8):210120. ISSN 2046-2441. doi: 10.1098/rsob.210120.

37. Kai Ma, Man Luo, Guanglei Xie, Xi Wang, Qilin Li, Lei Gao, Hongtao Yu, and Xiaochun Yu. Ribosomal RNA regulates chromosome clustering during mitosis. 8(1):1–13. ISSN 2056-5968. doi: 10.1038/s41421-022-00400-7. Publisher: Nature Publishing Group.

38. O. Lieleg, M. M. A. E. Claessens, Y. Luan, and A. R. Bausch. Transient binding and dissipation in cross-linked actin networks. 101(10):108101. doi: 10.1103/PhysRevLett.101.108101. Publisher: American Physical Society.

39. Norbert Willenbacher and Kristina Georgieva. Rheology of disperse systems. pages 7–49. Publisher: Wiley Online Library.

40. Je-Kyung Ryu, Allard J. Katan, Eli O. van der Sluis, Thomas Wisse, Ralph de Groot, Christian H. Haering, and Cees Dekker. The condensin holocomplex cycles dynamically between open and collapsed states. 27(12):1134–1141. ISSN 1545-9985. doi: 10.1038/s41594-020-0508-3. Publisher: Nature Publishing Group.

41. Oren Wintner, Nivi Hirsch-Attas, Miriam Schlossberg, Fani Brofman, Roy Friedman, Meital Kupervaser, Danny Kitsberg, and Amnon Buxboim. A unified linear viscoelastic model of the cell nucleus defines the mechanical contributions of lamins and chromatin. 7(8):1901222. ISSN 2198-3844. doi: 10.1002/advs.201901222. _eprint: https://onlinelibrary.wiley.com/doi/pdf/10.1002/advs.201901222.

42. Rosalia Ferraro, Stefano Guido, Sergio Caserta, and Manlio Tassieri. i-rheo-optical assay: Measuring the viscoelastic properties of multicellular spheroids. 26:101066. ISSN 2590-0064. doi: 10.1016/j.mtbio.2024.101066.

43. Timothy D. Matheson and Paul D. Kaufman. The p150n domain of chromatin assembly factor-1 regulates ki-67 accumulation on the mitotic perichromosomal layer. 28(1):21–29. ISSN 1059-1524. doi: 10.1091/mbc.e16-09-0659. Publisher: American Society for Cell Biology (mboc).

44. Kayo Hibino, Yuji Sakai, Sachiko Tamura, Masatoshi Takagi, Katsuhiko Minami, Toyoaki Natsume, Masa A. Shimazoe, Masato T. Kanemaki, Naoko Imamoto, and Kazuhiro Maeshima. Single-nucleosome imaging unveils that condensins and nucleosome–nucleosome interactions differentially constrain chromatin to organize mitotic chromosomes. 15(1):7152. ISSN 2041-1723. doi: 10.1038/s41467-024-51454-y. Publisher: Nature Publishing Group.

45. Karim Mrouj, Nuria Andrés-Sánchez, Geronimo Dubra, Priyanka Singh, Michal Sobecki, Dhanvantri Chahar, Emile Al Ghoul, Ana Bella Aznar, Susana Prieto, Nelly Pirot, Florence Bernex, Benoit Bordignon, Cedric Hassen-Khodja, Martin Villalba, Liliana Krasinska, and Daniel Fisher. Ki-67 regulates global gene expression and promotes sequential stages of carcinogenesis. 118(10):e2026507118. doi: 10.1073/pnas.2026507118. Publisher: Proceedings of the National Academy of Sciences.

46. Nuria Andrés-Sánchez, Daniel Fisher, and Liliana Krasinska. Physiological functions and roles in cancer of the proliferation marker ki-67. 135(11):jcs258932. ISSN 0021-9533. doi: 10.1242/jcs.258932.

47. C. Yang, J. Zhang, M. Ding, K. Xu, L. Li, L. Mao, and J. Zheng. Ki67 targeted strategies for cancer therapy. 20(5):570–575. ISSN 1699-3055. doi: 10.1007/s12094-017-1774-3.

48. Bin Wang, Lei Zhang, Tong Dai, Ziran Qin, Huasong Lu, Long Zhang, and Fangfang Zhou. Liquid–liquid phase separation in human health and diseases. 6(1):1–16. ISSN 2059-3635. doi: 10.1038/s41392-021-00678-1. Publisher: Nature Publishing Group.

49. Matthew G. Smith, Graham M. Gibson, and Manlio Tassieri. i-RheoFT: Fourier transforming sampled functions without artefacts. 11(1):24047. ISSN 2045-2322. doi: 10.1038/s41598-021-02922-8. Number: 1 Publisher: Nature Publishing Group.

50. Manlio Tassieri, Marco Laurati, Dan J. Curtis, Dietmar W. Auhl, Salvatore Coppola, Andrea Scalfati, Karl Hawkins, Phylip Rhodri Williams, and Jonathan M. Cooper. i-rheo: Measuring the materials’ linear viscoelastic properties “in a step “ ! 60(4):649–660. ISSN 0148-6055. doi: 10.1122/1.4953443. Publisher: Society of Rheology.

51. Curtis T Rueden, Johannes Schindelin, Mark C Hiner, Barry E DeZonia, Alison E Walter, Ellen T Arena, and Kevin W Eliceiri. ImageJ2: ImageJ for the next generation of scientific image data. 18(1):529. ISSN 1471-2105. doi: 10.1186/s12859-017-1934-z. Publisher: BioMed Central.

